# Highly divergent mesnidoviruses found in TSA database, especially in spiders

**DOI:** 10.1101/2025.01.04.631312

**Authors:** Jiazheng Xie, Yu Zhang

## Abstract

The mesnidoviruses are positive single-stranded viruses belonging to the order *Nidovirales*. They were mostly found in mosquitoes, and represent a link from small to large nidoviruses. In this study, we report 18 newly discovered nidoviruses in TSA database, most of them are from spiders (Order *Arachnida*). After reassembly, we got 9 nearly complete genomes with sizes ranged from ∼19,500 to 33,500 nt. Further phylogenetic analyses assigned these nidoviruses to *Mesnidovirineae*. Among them, the spider viruses represent a new clade. Pairwise evolutionary distance (PED) analysis showed these mesnidoviruses are highly divergent, with PED values from 2.2 to 4.9 with existing ones. Even among themselves the PED values are notable: with most exceeding the threshold of 0.32 that defines a new virus genus, suggesting these viruses are new species or belong to new higher-ranking taxa. In summary, we reported 18 mesnidoviruses in spiders, insects, plant and *Anthozoa*, the highly divergence suggested that spiders may harbor diverse mesnidoviruses. As the first nidovirus reported in *Arachnida* and *Anthozoa*, this discovery will contribute to the evolution study of *Nidovirales*.

---

The order *Nidovirales* exhibit a wide host range, with *Coronaviridae* (CoV), *Arteriviridae* (ArV) and *Torovirinae* (ToV) infecting vertebrate, *Roniviridae* (RoV) infecting crustaceans and *Mesnidovirineae* (MeV) infecting the insects. Among them, *Mesnidovirineae* contains two families: *Mesoniviridae* and *Medioniviridae*. Since their first discovery in mosquitoes in 2009 (1), mesoniviruses are thought to be mosquito specific. However, sporadic reports have emerged, indicating their presence in other species as well, such as aphid and thrip species (2, 3). They have attracted particular interest due to their intermediate genome sizes (20kb) between the small-sized nidoviruses (ArV, 13-16 kb) and the large-sized nidoviruses (CoV, ToV and RoV, 26-32 kb) (4, 5).

The organizations of mesnidovirus genomes are like other nidoviruses, with ORF1a and ORF1b occupying the 5’ two-thirds of its genome, the 3’-proximal one-third encoding viral structural proteins(6). The ORF1b overlaps with ORF1a, and its expression is regulated by ribosomal frame-shifting (RFS). In *Mesovirineae*, the RFS motif sequences are conserved with GGAUUUU (4, 7, 8). The 3’-proximal genes are transcribed from a nested set of 3’-coterminal sub-genomic (sg) mRNAs. These mRNAs co-terminate with 5’ leader sequences that are joined to the body transcription-regulating sequences (TRS).

Invertebrates harbor a diverse viruses (9). Among them, arthropods could play a central role in the evolution of viruses (10). Although it was hypothesized that *Nidovirales* may originated from arthropods (11), *Nidovirales* found in arthropods were limited to crustaceans and insects. As the second-largest class within arthropods, *Arachnida* encompass over 110,000 known species (12). However, viruses in *Arachnida* are greatly understudied, and no case of nidovirus in *Arachnida* has been reported.

With the development of sequencing technology, more and more studies and tools have been reported for detecting viruses within NGS data (13). For example, Serratus, a cloud computing infrastructure, has been specifically designed to enhance the detection of viruses in petabase-scale data (14), and LucaProt utilizes predicted structural information to search for viruses (15). Recently, numerous new vertebrate nidoviruses were discovered from searching public Sequence Read Archive database (16).

Here we first report highly divergent nidoviruses found in spider (*Arachnida*) and *Anthozoa* sequencing data in TSA database (17). Further phylogenetic analysis suggested that these viruses represent new mesnidoviruses. In summary, our study expanded the scale of the known host of nidoviruses and call for more research into nidoviruses in other species, especially in spiders (*Arachnida*).

## RESULTS

### Mesnidovirus contigs detected in TSA database

During our attempt to search coronaviruses in TSA database, a contig (GJFK01073640.1, *Stichodactyla gigantea*) aligned to the bovine coronavirus orf1ab with low identity 22.6%, but with long compared length (1120aa) was finally identified as mesnidovirus contig. Thus, we further searched the TSA database with mesnidovirus refseq sequences. Finally, 38 mesnidovirus contigs (Table S1) with identity ranging from 22.8% to 45.9% to existing mesnidoviruses were identified. Among them, 29 contigs were from 14 spider samples from bioproject PRJDB7399 (1000 spider silkomes) (18). 6 contigs were from two insects *Orius insidiosus* (19) and *Xestocephalus desertorum*, 1 contig from *Anthozoa Stichodactyla gigantea* (20), and 2 contigs from *Magnoliopsida* plant *Saccharum hybrid cultivar* (21).

### Reads depth and coverage

Following de novo genome assembly, we managed to retrieve 9 nearly complete genomes and 9 partial genomes. The length is between 6,324 to 33,469. We tentatively named these viruses following the pattern based on their host’s name: the first letter from the first word, then three letters from the second word, and finally the letter ‘V’ to signify that it is a virus (Table 1). Next, we aligned corresponding raw reads to the assembled genomes to calculate reads depth (Fig1 and Table S2). The results indicated that the proportion of reads belonging to mesnidoviruses in the raw data can be high, such as 4.6% for C.uniV (*Cheiracanthium unicum mesnidovirus*), 5.26% for L.minV (*Lepthyphantes minutus mesnidovirus*) and 11.3% for M.pseV (*Microdipoena pseudojobi mesnidovirus*). Given that the initial purpose of these spider transcriptome data was to study silk gene in spiders (18), not for viral identification, the observed high viral reads proportion suggests that these field-collected spider samples were naturally infected with mesnidoviruses. The average depths are uneven across samples, from 14 x to 15,610 x (L.minV). What’s more, corresponding to the functional regions encoding structural proteins of coronaviruses, the reads depth is significantly higher at the 3’ end of their genomes than in other regions.

**Table 1.** Assembled genomes of new mesnidoviruses.

| Virus abbreviation | TSA Index | Sample species name | Scaffold Length |
| --- | --- | --- | --- |
| S.gigV | GJFK01 | *Stichodactyla gigantea* | 10619 |
| P.opiV | IALF01 | *Pholcus opilionoides* | 13157 |
| U.comV | IAMC01 | *Uroctea compactilis* | 27241 |
| H.troV | IAOO01 | *Hickmania troglodytes* | 7733 |
| T.genIDV6716V | IAPB01 | *Theridiidae gen. sp. IDV 6716* | 17412 |
| T.genIDV5269V | IBEI01 | *Theridiidae gen. sp. IDV 5269* | 33469 |
| M.tokV | IBFE01 | *Mecopisthes tokumotoi* | 15851 |
| A.yagV | IBNT01 | *Araniella yaginumai* | 29222 |
| A.stuV | IBWJ01 | *Anelosimus studiosus* | 15245 |
| E.morV | IBXH01 | *Eutittha mordax* | 16382 |
| L.minV | IBYE01 | *Lepthyphantes minutus* | 28652 |
| X.desV | GELC01 | *Xestocephalus desertorum* | 22527 |
| S.hybV | GFHZ01 | *Saccharum hybrid cultivar* | 20381 |
| O.insV | GCYL01 | *Orius insidiosus* | 19436 |
| N.varV | IAIY01 | *Nerienne variabilis* | 6324 |
| C.uniV | IAWR01 | *Cheiracanthium unicum* | 30492 |
| M.pseV | IBIF01 | *Microdipoena pseudojobi* | 31892 |
| H.mitV | ICNE01 | *Hentzia mitrata* | 6590 |

**Figure 1.**
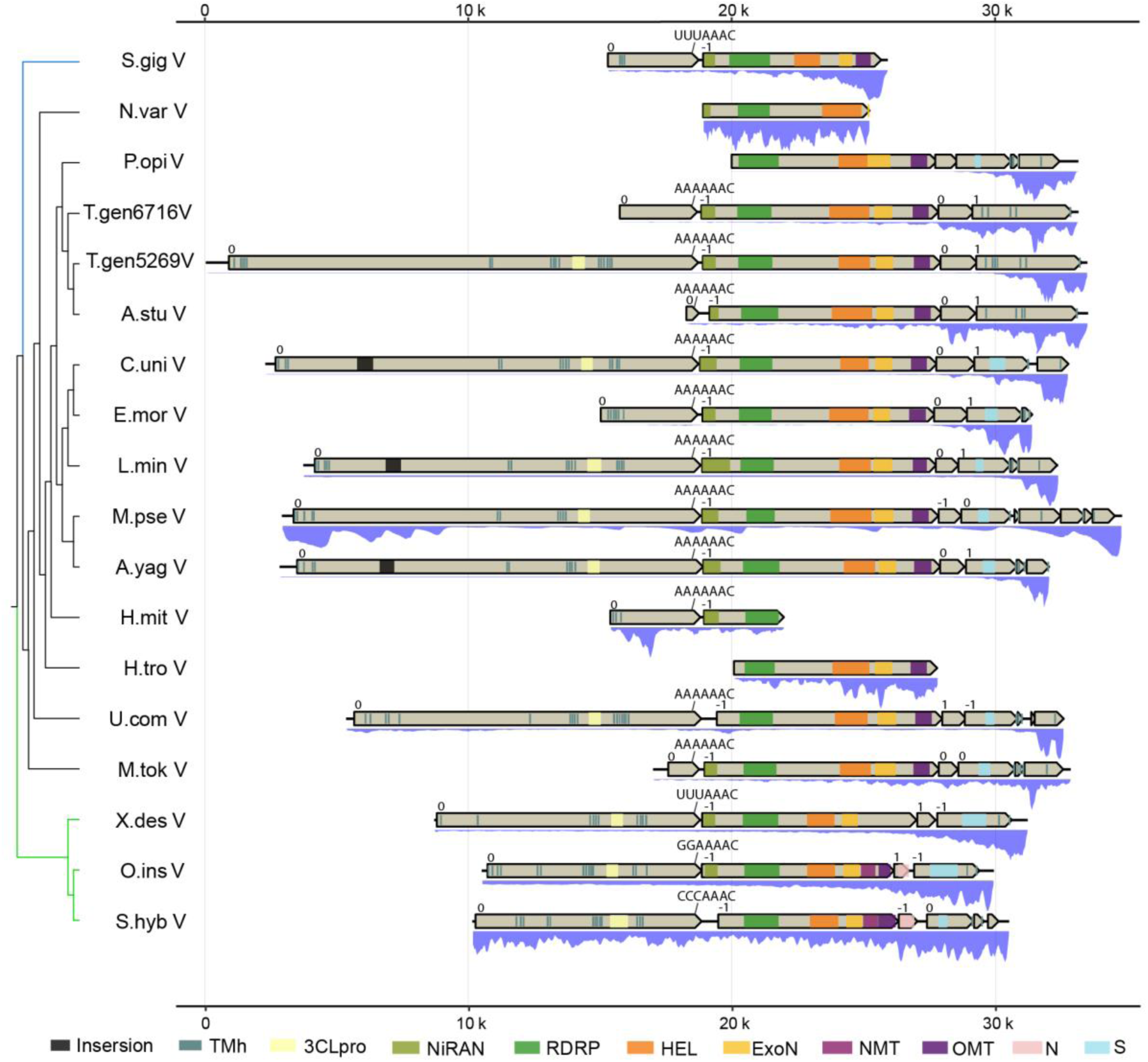
Genome organization of newly found mesnidoviruses. Open reading frames are showed as rectangles with arrows. Protein domains, and sequence insertions are indicated in colored box; transmembrane helix (TMh), 3C-like protease (3CLpro), RdRp-associated nucleotidyltransferase (NiRAN), RNA-dependent RNA polymerase (RdRp), superfamily 1 helicase (Hel), 3’-5’ exoribonuclease (ExoN), Guanine-N7methyltransferase (NMT), Omethyltransferase (OMT), nucleocapsid protein (N), spike (S). The depth plots below the genome are generated by normalizing the depth of each site, dividing by the maximum depth observed in each genome.

### Genome organization

The organizations of these mesnidovirus genomes were similar to other nidoviruses (Fig1), featuring approximately six major open reading frames (ORFs), with ORF1a and ORF1b occupying the 5’ four-fifths of the genome. The length of ORF1a are different among different viruses, it is the main reason that these viruses have different length. The 3CLpro domain and the transmembrane domains flanking 3CLpro are conserved. The ORF1b overlaps ORF1a in the −1 frame. Using the pattern (XXXYYYZ) conserved in vertebrate nidoviruses to search these mesnidovirus genomes, we found a heptamer AAAAAAC about 40 nt upstream of ORF1a stop codon in all spider mesnidoviruses. AAAAAAC is also the frame shift sequence used by the astrovirus (22). As for viruses from other species, UUUAAAC was found for S.gigV (*Stichodactyla gigantea mesnidovirus*) and X.desV (*Xestocephalus desertorum mesnidovirus*), CCCAAAC for S.hybV (*Saccharum hybrid cultivar mesnidovirus*). However, we did not find the XXXYYYZ pattern as anticipated for O.insV (*Orius insidiosus mesnidovirus*); instead, a GGAAAAC most likely was identified 19bp upstream of the ORF1a stop codon. Although 3’-5’ exoribonuclease (ExoN) and 2’-O-methyltransferase (OMT) are also conserved across most of these mesnidoviruses, Guanine-N7methyltransferase (NMT) domain is less conserved. Only in the insect/plant mesnidoviruses (O.insV and S.hybV), NMT domain was found by Interproscan. Notably, between ∼1000-1200 of orf1a in three spider viruses (C.uniV, L.minV, A.yagV (*Araniella yaginumai mesnidovirus*)), there was a ∼200aa insertion with highest identity to spider encoded RING-type domain-containing protein. This may reflect the complex interplay resulting from long-term virus-host associations.

### Structure proteins

By contrast, the structure proteins of nidoviruses are less conserved. Using structure proteins of existing nidoviruses to searching similarity with our newly discovered mesnidoviruses, only nucleotide capsid (N) and spike protein (S) proteins exhibit gene homology in some of these mesnidoviruses. For example, the N proteins are conserved in the insect/plant mesoniviruses (O.insV and S.hybV). While in three spider mesnidoviruses, T.gen 6716V (*Theridiidae gen. sp. IDV 6716 mesnidovirus*), T.gen 5269V (*Theridiidae gen. sp. IDV 5269 mesnidovirus*) and A.stuV (*Anelosimus studiosus mesnidovirus*), S protein homology can’t be identified. However, in these three viruses the orf4 (∼1250aa) exhibit the highest identities (∼20%) with another virus identified in spider (YP_009336501.1, *Hubei tetragnatha maxillosa virus 7*) (Table S3). Viruses frequently swap functional components, with a notable propensity for the transfer of structural proteins that can span significant evolutionary gaps (9). Thus, the S protein in T.gen 6716V, T.gen 5269V and A.stuV may have undergone recombination with the analogous protein from another spider virus *Hubei tetragnatha maxillosa virus 7*.

### Phylogenetic analysis

Next, we proceeded to analyze the phylogenetic position of these new mesnidoviruses. Conserved domains (RdRp, Hel and ExoN) of these mesnidoviruses were concatenated, and used for phylogenetic tree inferring. The best model is LG+F+R6 according Bayesian Information Criterion (BIC). The phylogenetic tree showed viruses found in this study are located in the mesnidoviruses clade (Fig2A), with two insect viruses (O.insV and X.desV) and the plant virus (S.hybV) located in mesoniviruses, the O.insV is closest to mesonivirus found in aphis insect. The Anthozoa virus (S.gigV) was located in *Medioniviridae*. Interestingly, X.desV is located between mesoniviruses and *Medioniviridae*. 14 mesnidoviruses from spiders formed a separate clade. As some genomes are incomplete, we further inferred a root tree using all RdRp domains of these viruses (Fig2B). Result is the same: these newly found spider mesnidovirus formed a new clade beside *Medioniviridae*.

**Figure 2.**
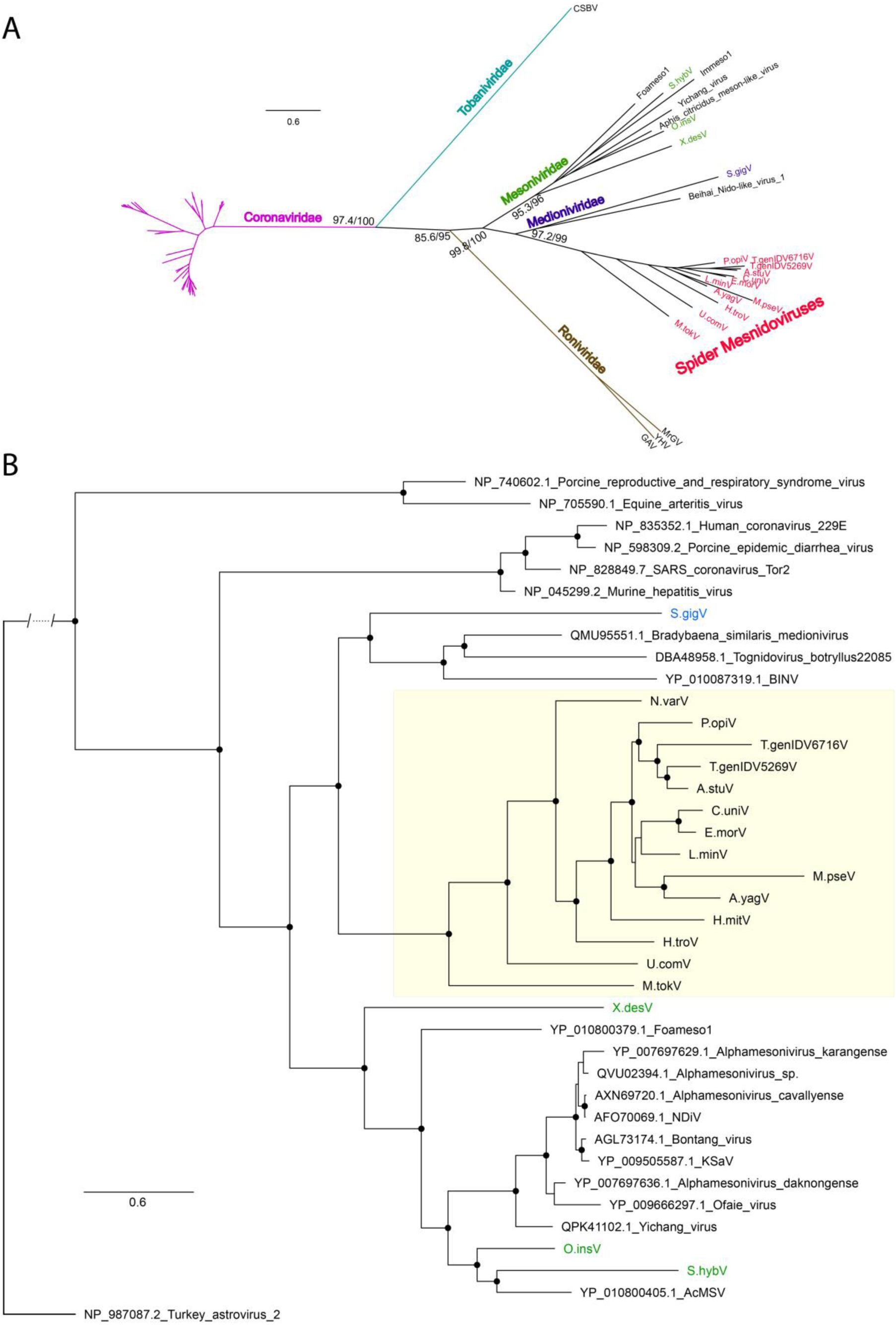
Phylogram for newly discovered mesnidoviruses. (A) Unrooted tree inferred based on alignment of concatenated conserved domain RdRp, Hel and ExoN. Branch support values are showed with Shimodaira-Hasegawa Likelihood Ratio Test (SH-aLRT) support / bootstrap percentage from 1000 replicate ultrafast bootstrap searches. (B) Root tree based on RdRp domain and including astrovirus as outgroup.

To verify whether these viruses are new viruses, we calculated the PED values with the 9 nearly full genome using concatenated domains of 3CLpro, RdRp, Hel, ExoN (Table S4). The PED values showed that all the 9 mesnidoviruses have PED greater than 0.32 (threshold recognized for genus separation). Therefore, these viruses likely belong to an unassigned genus. As some virus genomes are not complete, we also calculated the PED with RdRp alone, and found among the 18 mesnidoviruses, only 4 viruses have paired PED less than 0.32. They are E.morV *VS.* C.uniV (0.21) and T.genIDV5269V *VS.* A.stuV. What’s more, using more domains like concatenated 3CLpro, Hel, RdRp, ExoN will generate larger PED values because other domains are less conserved than RdRp. Therefore, these 18 viruses are all new mesnidoviruses.

### Highly divergent 3CLpro but key functional residues conserved

As in these viruses, 3CLpro domains cannot been predicted by Interproscan except in two insect viruses (O.insV and X.desV), we speculated that 3CLpro are highly divergent in these viruses. To determine whether 3CLpro are functional, we used blast to identified 3CLpro domains of these viruses, and performed multiple sequence alignments. The result showed: 3CLpro are highly divergent, but the key residues are conserved (Fig 3). The catalytic Cys (1539) -His (1434) dyad are conserved in all these 18 viruses. Apart from that, H1530, T1534, H1554 are also highly conserved. While the functions of T1534, H1554 have been deduced (23), highly conserved H1530 remains unexplored.

**Figure 3.**
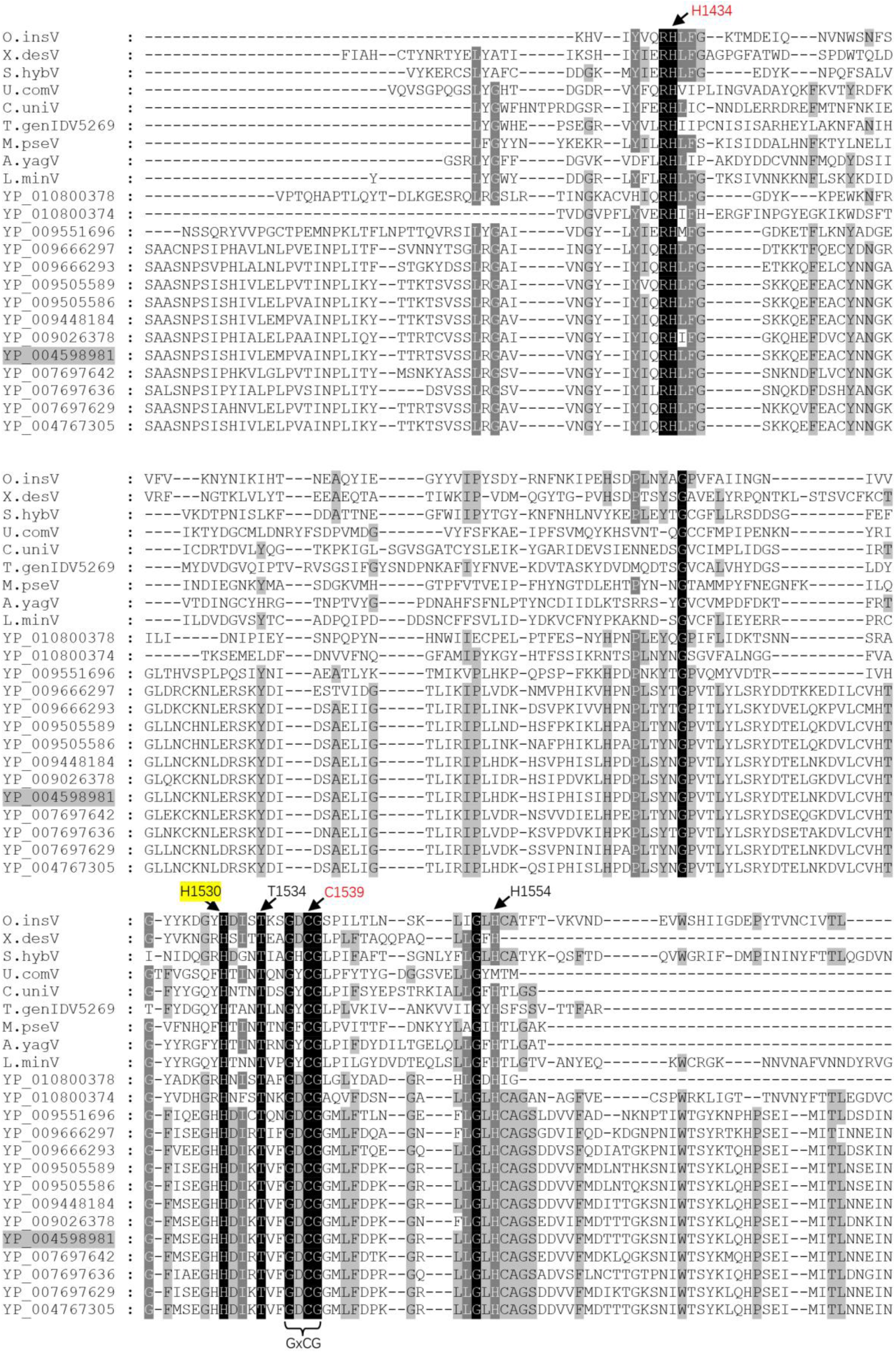
Sequence alignment of newly discovered mesnidovirus 3CLpro domains. Show are the putative 3CLpro domains of newly discovered and representative mesnidoviruses, along with their functional residues. Amino acid numbering corresponds to the position in the amino acid sequence of CavV pp1ab (YP_004598981).

### Transcription regulatory sequences

At last, we speculate the TRS by searching leader-body junction reads that span ORF1a1b, and mapping both 5’ leading sequence and TRS located immediately upstream of the structure proteins. We found different viruses used different TRS (Table 2 & FigS1). Such as C?TCGTCTC(T|G)ATT(T|C)T(T|C)AGA found in T.genIDV5269V and AACTT(A)AGCAT(TC)C in L.minV.

**Table 2.** Putative transcription-regulating sequences (TRS) deduced from the reads crossing the junctions between 5’ leader sequence and the TRS upstream of structure proteins.

| Viruses | Putative TRS | Genome position |
| --- | --- | --- |
| T.genIDV5269V | TCGTCTC(T\|G)ATT(T\|C)T(T\|C)AGA | 152 |
|  |  | 27,837 |
|  |  | 28,896 |
| L.minV | AACT(T\|C)(C\|A)AGCA | 74 |
|  |  | 23,973 |
|  |  | 27,014 |
|  |  | 28,575 |

## DISCUSSION

In this study, we described discovery of 18 new mesnidoviruses in TSA database. After assembly, 9 nearly complete genomes were got. Among them, nidoviruses found in Arachnida and Anthozoa are first time reported. PED and phylogenetic analysis showed that these mesnidoviruses are highly divergent. However, the functionally critical residues of functional domains are preserved. Notably, spider host sequence integration was observed at the 5’ of ORF1a and similarity found between the structure protein and another spider virus in some spider mesnidoviruses. These observations suggested mesnidoviruses have long co-existed with spiders. And further studies of spider viruses are needed to uncover the nature of highly divergent nidoviruses.

Another interesting question is that why mesnidoviruses showed such a high divergence. In this study, 18 mesnidoviruses were found, with only two paired PED values less than 0.32 using RdRp alone. As RdRp showed a maximum sequence identity, the PED using more domains would be larger. ExoN is the enzyme that can correct the wrong base to assure the replication fidelity. However, ExoNs are all conserved in these mesnidoviruses. Thus, the underlying reason for the high divergence needs to be further invested by future research efforts.

Insects can transmit plant viruses while they are feeding on injected plants, vice versa. Thus, mesnidoviruses found in the plant in this study (S.hybV) would be actually be insect originated.

In summary, besides Roniviridae in crustaceans, and mesoniviruses in insects, in this stury, our discovery of mesnidoviruses in spiders further expanded the host range of nidoviruses to the second largest clade of *Arthropoda*: *Arachnida*. As mesnidoviruses reported are most in the intermediate genome size (20,000nt), our newly found mesnidoviruses (19,500-33,500nt) also significantly expands the known length range of mesnidoviruses.

## MATERIALS AND METHODS

### Mesnidoviruses searching in TSA database

TSA data were downloaded from Genbank (https://www.ncbi.nlm.nih.gov/genbank/tsa/). Refseq protein sequences of mesnidoviruses (taxid 2499400) were formated as reference database and aligned to TSA contigs with --evalue 1e-05 using Diamond (24). The contigs hit were further blasted to core-nt database to remove false positives that aligned to non-virus sequences.

### Virus genome assembly

After viral contigs identified, we downloaded the corresponding raw data from the Sequence Read Archive (SRA). Trimmomatic was used to filter low-quality reads (25). Virus genomes were assembled by MEGAHIT (26). Resulting scaffolds were further blasted to mesnidovirus database to remove non-mesnidovirus sequences. After virus genomes determined, reads were aligned to the genomes with hisat2 and visualized by IGV-tools (27, 28). The alignment SAM files were dealt with samtools and depth calculated with bedtools (29, 30).

### Protein domain and phylogenetic analysis

Open reading frames (ORFs) were predicted with prodigal (31). Domains were annotated with Interproscan (32), and transmebrane domains were predicted with tmhmm2 (33). Conserved domains of RNA polymerases (RdRp), helicase (Hel), 3’-5’ exoribonuclease (ExoN) were aligned with MAFFT (34). The maximum likelihood (ML) Phylogenetic tree and pairwise evolutionary distances (PED) were inferred with IQ-tree with 1000 replicates for ultrafast bootstrap and 1000 replicates for SH approximate likelihood ratio test (35). Finally, the phylogenetic tree was edit by Figtree V1.44 (https://github.com/rambaut/figtree/).

## SUPPLEMENTAL INFORMATION

Supplemental Information includes one figure, and six tables.

## AUTHOR CONTRIBUTIONS

J.Z.X, conceived the concept, J.Z.X and Y.Z performed the bioinformatic analysis. J.Z.X, Y.Z prepared the draft.

## ACKNOWLEDGEMENTS

We express our gratitude to all colleagues within the scientific community for making their sequencing data publicly accessible. We also acknowledge the National Center for Biotechnology Information (NCBI) for providing a comprehensive platform for the exchange of sequencing data. This work was supported by Natural Science Foundation of China (Grant No. 31900152) and Chongqing Science and Technology Bureau (Grant No. CSTB2024NSCQ-MSX1213).

## EXTENDED DATA LEGENDS

**Figure 1.**
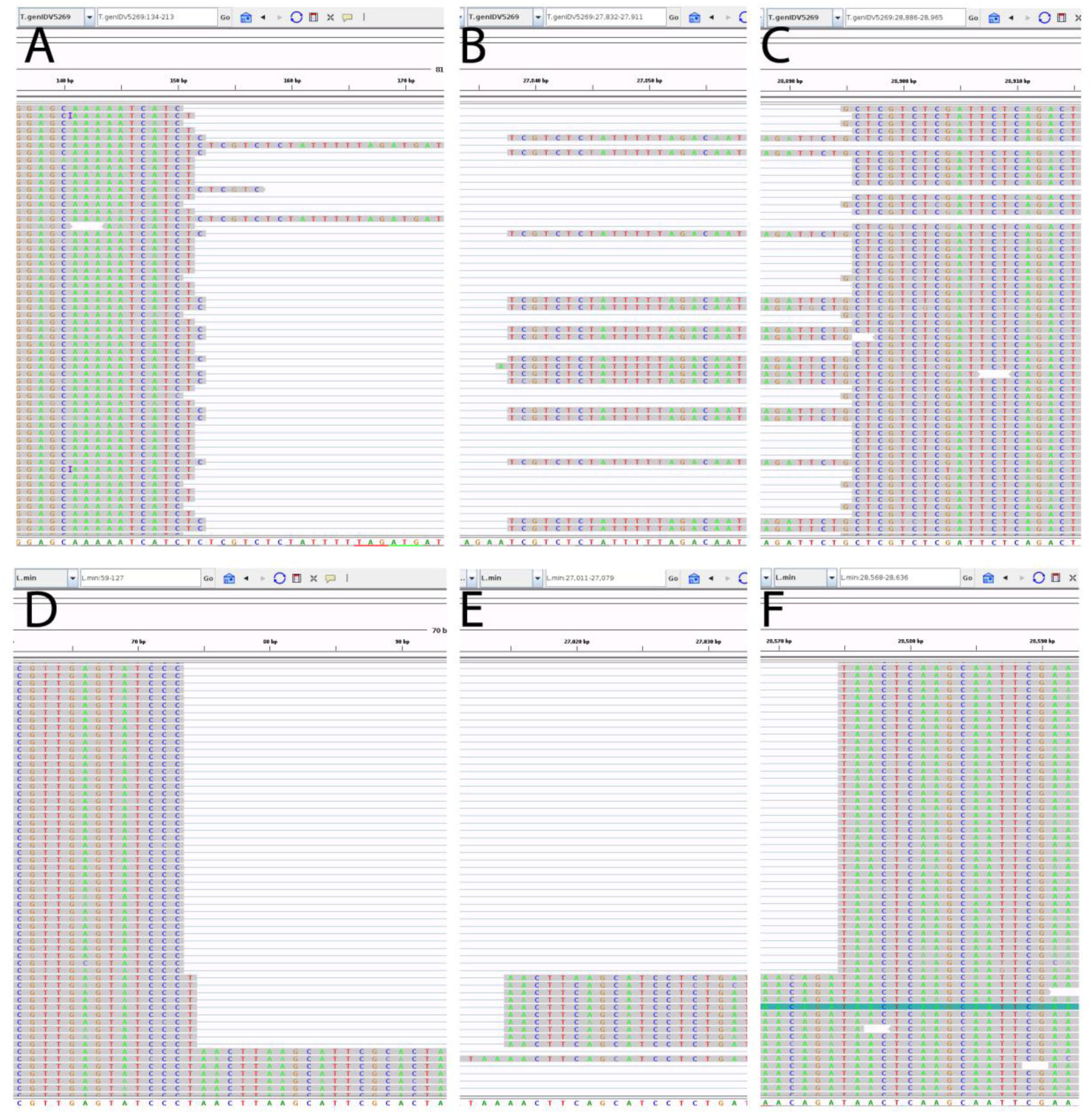
IGV views of reads crossed the ORF1ab junction that mapped to both leader transcription regulatory sequence (TRS-L) and the body transcription regulatory sequences (TRS-Bs). (A) Screenshot of the 5’ part of reads across regions of 152 to 27,837/28,896 in T.genIDV5269V. (B) Screenshot of the 3’ part of reads across regions of 152 to 27,837 in T.genIDV5269V. (C) Screenshot of the 3’ part of reads across regions of 152 to 28,896 in T.genIDV5269V. (D) Screenshot of the 5’ part of reads across regions of 74 to 27,014/28,575 in L.minV. (E) Screenshot of the 3’ part of reads across regions of 74 to 27,014 in L.minV. (F) Screenshot of the 3’ part of reads across regions of 74 to 28,575 in L.minV.

**Extended Data Table 1.** List of contigs that mapped to mesnidoviruses.

| Contigs | Biosample | Biosample source species name | Biosample source Class | Mapped reference | Mapped reference protein description | Mapped identity | Mapped length | Evalue |
| --- | --- | --- | --- | --- | --- | --- | --- | --- |
| GFHZ01020763.1 | SAMN06323325 | *Saccharum hybrid cultivar* | Magnoliopsida | QLL27750.1 | putative peptidase, partial [Leveillula taurica associated alphamesonivirus 1] | 29.4 | 1103 | 3.44E-100 |
| GFHZ01115333.1 | SAMN06323325 | *Saccharum hybrid cultivar* | Magnoliopsida | QLL27747.1 | putative helicase, partial [Leveillula taurica associated alphamesonivirus 1] | 36.5 | 1239 | 1.05E-225 |
| GCYL01017727.1 | SAMN03341986 | *Orius insidiosus* | Insecta | YP_010800405.1 | putative ORF1ab [Aphis citricidus meson-like virus] | 31.4 | 1676 | 1.21E-168 |
| GCYL01017728.1 | SAMN03341986 | *Orius insidiosus* | Insecta | YP_010800408.1 | putative ORF2a [Aphis citricidus meson-like virus] | 25.6 | 418 | 3.63E-21 |
| GCYL01017729.1 | SAMN03341986 | *Orius insidiosus* | Insecta | YP_010800405.1 | putative ORF1ab [Aphis citricidus meson-like virus] | 45.9 | 632 | 2.63E-147 |
| GCYL01017730.1 | SAMN03341986 | *Orius insidiosus* | Insecta | YP_010800405.1 | putative ORF1ab [Aphis citricidus meson-like virus] | 39 | 420 | 1.29E-76 |
| GCYL01017731.1 | SAMN03341986 | *Orius insidiosus* | Insecta | QPK41102.1 | putative replicase polyprotein [Yichang virus] | 39.9 | 203 | 2.30E-30 |
| GELC01073079.1 | SAMN04101404 | *Xestocephalus desertorum* | Insecta | YP_010800405.1 | putative ORF1ab [Aphis citricidus meson-like virus] | 29.1 | 2196 | 1.95E-220 |
| GJFK01073640.1 | SAMN18817344 | *Stichodactyla gigantea* | Anthozoa | YP_009333345.1 | hypothetical protein [Beihai Nido-like virus 1] | 25.3 | 2418 | 1.87E-139 |
| IALF01011102.1 | SAMD00139091 | *Pholcus opilionoides* | Arachnida | YP_009333345.1 | hypothetical protein [Beihai Nido-like virus 1] | 24.1 | 1458 | 1.32E-61 |
| IAMC01016640.1 | SAMD00139114 | *Uroctea compactilis* | Arachnida | YP_010087319.1 | orf1ab polyprotein [Botryllodes leachii nidovirus] | 26.6 | 979 | 2.77E-77 |
| IAMC01018833.1 | SAMD00139114 | *Uroctea compactilis* | Arachnida | YP_010087319.1 | orf1ab polyprotein [Botryllodes leachii nidovirus] | 26.6 | 979 | 3.62E-77 |
| IAMC01021898.1 | SAMD00139114 | *Uroctea compactilis* | Arachnida | YP_010087319.1 | orf1ab polyprotein [Botryllodes leachii nidovirus] | 26.6 | 979 | 2.78E-77 |
| IAMC01022801.1 | SAMD00139114 | *Uroctea compactilis* | Arachnida | YP_010087319.1 | orf1ab polyprotein [Botryllodes leachii nidovirus] | 26.6 | 979 | 3.63E-77 |
| IAMC01036124.1 | SAMD00139114 | *Uroctea compactilis* | Arachnida | YP_010087319.1 | orf1ab polyprotein [Botryllodes leachii nidovirus] | 26.6 | 979 | 3.61E-77 |
| IAOO01004986.1 | SAMD00139178 | *Hickmania troglodytes* | Arachnida | YP_007697638.1 | ORF2a [Alphamesonivirus daknongense] | 43.1 | 51 | 7.31E-06 |
| IAOO01036033.1 | SAMD00139178 | *Hickmania troglodytes* | Arachnida | YP_010800405.1 | putative ORF1ab [Aphis citricidus meson-like virus] | 23.8 | 1490 | 2.41E-51 |
| IAPB01015957.1 | SAMD00139191 | *Theridiidae gen. sp. IDV 6716* | Arachnida | YP_009666293.1 | putative ORF1ab [Kadiweu virus] | 26.1 | 1101 | 1.17E-78 |
| IBEI01031789.1 | SAMD00139588 | *Theridiidae gen. sp. IDV 5269* | Arachnida | YP_010800405.1 | putative ORF1ab [Aphis citricidus meson-like virus] | 23.8 | 1340 | 7.35E-31 |
| IBEI01037011.1 | SAMD00139588 | *Theridiidae gen. sp. IDV 5269* | Arachnida | YP_009666293.1 | putative ORF1ab [Kadiweu virus] | 26.2 | 1106 | 4.92E-78 |
| IBFE01009263.1 | SAMD00139610 | *Mecopisthes tokumotoi* | Arachnida | YP_009333345.1 | hypothetical protein [Beihai Nido-like virus 1] | 26.7 | 967 | 5.25E-75 |
| IBNT01009445.1 | SAMD00139833 | *Araniella yaginumai* | Arachnida | YP_010800405.1 | putative ORF1ab [Aphis citricidus meson-like virus] | 26.9 | 949 | 9.45E-73 |
| IBNT01011284.1 | SAMD00139833 | *Araniella yaginumai* | Arachnida | YP_010800405.1 | putative ORF1ab [Aphis citricidus meson-like virus] | 26.9 | 949 | 9.63E-73 |
| IBNT01028316.1 | SAMD00139833 | *Araniella yaginumai* | Arachnida | YP_009505591.1 | putative spike glycoprotein [Dak Nong virus] | 33.6 | 134 | 5.95E-10 |
| IBWJ01020797.1 | SAMD00140057 | *Anelosimus studiosus* | Arachnida | YP_010087319.1 | orf1ab polyprotein [Botryllodes leachii nidovirus] | 28.4 | 377 | 3.26E-26 |
| IBWJ01031957.1 | SAMD00140057 | *Anelosimus studiosus* | Arachnida | YP_009333345.1 | hypothetical protein [Beihai Nido-like virus 1] | 22.8 | 1055 | 8.65E-39 |
| IBWJ01039700.1 | SAMD00140057 | *Anelosimus studiosus* | Arachnida | YP_009333345.1 | hypothetical protein [Beihai Nido-like virus 1] | 34.9 | 298 | 1.58E-35 |
| IBXH01002215.1 | SAMD00140081 | *Eutitha mordax* | Arachnida | YP_010087319.1 | orf1ab polyprotein [Botryllodes leachii nidovirus] | 25.8 | 1055 | 1.35E-78 |
| IBXH01019418.1 | SAMD00140081 | *Eutitha mordax* | Arachnida | YP_010087319.1 | orf1ab polyprotein [Botryllodes leachii nidovirus] | 25.8 | 1055 | 1.34E-78 |
| IBYE01022616.1 | SAMD00140104 | *Lepthyphantes minutus* | Arachnida | YP_009666293.1 | putative ORF1ab [Kadiweu virus] | 27.4 | 1099 | 1.23E-81 |
| IBYE01030818.1 | SAMD00140104 | *Lepthyphantes minutus* | Arachnida | YP_009666293.1 | putative ORF1ab [Kadiweu virus] | 27.4 | 1099 | 1.25E-81 |
| IAIY01003509.1 | SAMD00139032 | *Neriene variabilis* | Arachnida | AGE00056.1 | ORF1ab [Hana virus] | 27.2 | 952 | 2.01E-71 |
| IAWR01001386.1 | SAMD00139389 | *Cheiracanthium unicum* | Arachnida | YP_007697644.1 | ORF2a [Alphamesonivirus casuarinaense] | 26.5 | 151 | 1.41E-09 |
| IAWR01017767.1 | SAMD00139389 | *Cheiracanthium unicum* | Arachnida | YP_009666293.1 | putative ORF1ab [Kadiweu virus] | 27.3 | 1094 | 3.99E-78 |
| IAWR01036719.1 | SAMD00139389 | *Cheiracanthium unicum* | Arachnida | YP_009666293.1 | putative ORF1ab [Kadiweu virus] | 27.3 | 1094 | 4.04E-78 |
| IBIF01008746.1 | SAMD00139689 | *Microdipoena pseudojovi* | Arachnida | YP_010087319.1 | orf1ab polyprotein [Botryllodes leachii nidovirus] | 24.9 | 1432 | 5.97E-61 |
| IBIF01015685.1 | SAMD00139689 | *Microdipoena pseudojovi* | Arachnida | YP_009666293.1 | putative ORF1ab [Kadiweu virus] | 25.3 | 1088 | 1.84E-63 |
| ICNE01003722.1 | SAMD00157222 | *Hentzia mitrata* | Arachnida | YP_009666293.1 | putative ORF1ab [Kadiweu virus] | 26.3 | 1125 | 1.05E-76 |

**Extended Data Table 2.** The percentage and depth of reads mapped to the newly discovered mesnidoviruses.

| Mesnidoviruses | TSA Index | Sample species name | Scaffold Length | Mapped reads number | Total reads number | Percentage of total reads(%) | Average depth |
| --- | --- | --- | --- | --- | --- | --- | --- |
| S.gigV | GJFK01 | *Stichodactyla gigantea* | 10619 | 2785 | 80697418 | 0.003451164 | 51 |
| P.opiV | IALF01 | *Pholcus opilionoides* | 13157 | 191302 | 21306081 | 0.897875118 | 2798 |
| U.comV | IAMC01 | *Uroctea compactilis* | 27241 | 358912 | 14242187 | 2.520062403 | 2809 |
| H.troV | IAOO01 | *Hickmania troglodytes* | 7733 | 1713 | 19686064 | 0.008701587 | 41 |
| T.genIDV6716V | IAPB01 | *Theridiidae* gen. sp. IDV 6716 | 17412 | 46023 | 20673129 | 0.222622323 | 510 |
| T.genIDV5269V | IBEI01 | *Theridiidae* gen. sp. IDV 5269 | 33469 | 374546 | 18659233 | 2.007295798 | 2561 |
| M.tokV | IBFE01 | *Mecopisthes tokumotoi* | 15851 | 13269 | 21831290 | 0.060779734 | 156 |
| A.yagV | IBNT01 | *Araniella yaginumai* | 29222 | 121318 | 13957111 | 0.869219998 | 1130 |
| A.stuV | IBWJ01 | *Anelosimus studiosus* | 15245 | 9456 | 19215760 | 0.049209607 | 120 |
| E.morV | IBXH01 | *Eutitha mordax* | 16382 | 51824 | 20308784 | 0.255180222 | 605 |
| L.minV | IBYE01 | *Leptyphantes minutus* | 28652 | 901278 | 17138989 | 5.258641569 | 15610 |
| X.desV | GELC01 | *Xestocephalus desertorum* | 22527 | 24582 | 25906961 | 0.094885695 | 177 |
| S.hybV | GFHZ01 | *Saccharum hybrid cultivar* | 20381 | 4768 | 754933543 | 0.000631579 | 31 |
| O.insV | GCYL01 | *Orius insidiosus* | 19436 | 48824 | 24441305 | 0.199760201 | 566 |
| N.varV | IAIY01 | *Nerene variabilis* | 6324 | 478 | 18528530 | 0.002579805 | 14 |
| C.uniV | IAWR01 | *Cheiracanthium unicum* | 30492 | 827580 | 17950781 | 4.610272946 | 6502 |
| M.pseV | IBIF01 | *Microdipoena pseudojobi* | 31892 | 1525092 | 13447121 | 11.34140163 | 9397 |
| H.mitV | ICNE01 | *Hentzia mitrata* | 6590 | 1426 | 21755810 | 0.006554571 | 41 |

**Extended Data Table 3.** List of top score hits of insersion segment (in orf1a) or structure proteins (orf4/5/6) blasted to nr database.

| query | subject | identity | Length<br>(aa) | mismatch | Gap | start | end | Subject<br>start | subend | evalue | score | subject name | subject class |
| --- | --- | --- | --- | --- | --- | --- | --- | --- | --- | --- | --- | --- | --- |
| C.uniV<br>(orf1a) | GFR32553.1 | 30.275 | 218 | 107 | 6 | 1023 | 1224 | 34 | 222 | 7.01E-15 | 90.9 | Trichonephila clavata | Arachnida |
| C.uniV<br>(orf1a) | GFT61753.1 | 30.909 | 220 | 105 | 8 | 1022 | 1224 | 33 | 222 | 4.74E-12 | 82.4 | Trichonephila clavipes | Arachnida |
| C.uniV<br>(orf1a) | GBN88177.1 | 31.441 | 229 | 118 | 9 | 1011 | 1224 | 27 | 231 | 8.40E-11 | 79.7 | Araneus ventricosus | Arachnida |
| L.minV<br>(orf1a) | GBN16693.1 | 28.638 | 213 | 114 | 8 | 888 | 1082 | 29 | 221 | 2.20E-11 | 80.5 | Araneus ventricosus | Arachnida |
| L.minV<br>(orf1a) | GBN88177.1 | 26.568 | 271 | 155 | 9 | 888 | 1140 | 38 | 282 | 7.56E-11 | 79.7 | Araneus ventricosus | Arachnida |
| L.minV<br>(orf1a) | GFT61753.1 | 32.02 | 203 | 99 | 8 | 893 | 1078 | 39 | 219 | 2.87E-10 | 77 | Trichonephila clavipes | Arachnida |
| A.yagV<br>(orf1a) | GFR32553.1 | 30.594 | 219 | 139 | 4 | 1053 | 1264 | 40 | 252 | 3.78E-20 | 105 | Trichonephila clavata | Arachnida |
| A.yagV<br>(orf1a) | GBN88177.1 | 28.033 | 239 | 148 | 7 | 1049 | 1269 | 40 | 272 | 1.66E-17 | 99.8 | Araneus ventricosus | Arachnida |
| A.yagV<br>(orf1a) | GFT61753.1 | 30.045 | 223 | 135 | 6 | 1053 | 1264 | 40 | 252 | 1.20E-16 | 95.9 | Trichonephila clavipes | Arachnida |
| S.hybV<br>(orf6) | QQG34660.1 | 39.362 | 94 | 56 | 1 | 24 | 117 | 45 | 137 | 2.55E-08 | 60.8 | Nasturtium officinale<br>macula-like virus 1 | Alsuviricetes |
| T.genIDV<br>6716V<br>(orf4) | YP_009336501.1 | 21.931 | 839 | 556 | 26 | 403 | 1227 | 942 | 1695 | 2.02E-33 | 155 | Hubei tetragnatha<br>maxillosa virus 7 | Arachnida<br>Riboviria |
| T.genIDV<br>5269V<br>(orf5) | YP_009336501.1 | 23.443 | 819 | 512 | 31 | 528 | 1312 | 983 | 1720 | 2.96E-38 | 171 | Hubei tetragnatha<br>maxillosa virus 7 | Arachnida<br>Riboviria |
| A.stuV<br>(orf4) | YP_009336501.1 | 21.787 | 817 | 512 | 32 | 473 | 1261 | 983 | 1700 | 1.52E-25 | 129 | Hubei tetragnatha<br>maxillosa virus 7 | Arachnida<br>Riboviria |

**Extended Data Table 4.** Pairwise evolutionary distance (PED) values using conserved domains (3CLpro, RdRp, Hel and ExoN).

|  | O.insV | X.desV | S.hybV | U.comV | C.uniV | T.genIDV<br>5269V | M.pseV | A.yagV | L.minV | Foameso1 | Immeso1 | YCV | OFAV | KADV | DKNV | KSaV | DKV | CASV | CavV | NDiV |
| --- | --- | --- | --- | --- | --- | --- | --- | --- | --- | --- | --- | --- | --- | --- | --- | --- | --- | --- | --- | --- |
| O.insV | 0.00 | 2.70 | 2.39 | 4.56 | 4.24 | 4.49 | 4.30 | 4.35 | 4.21 | 2.39 | 2.41 | 2.23 | 2.25 | 2.31 | 2.28 | 2.29 | 2.35 | 2.32 | 2.29 | 2.28 |
| X.desV | 2.70 | 0.00 | 3.22 | 4.39 | 4.25 | 4.37 | 4.29 | 4.59 | 4.30 | 2.83 | 3.28 | 3.04 | 3.18 | 3.06 | 2.94 | 2.94 | 2.96 | 2.97 | 2.96 | 2.95 |
| S.hybV | 2.39 | 3.22 | 0.00 | 4.49 | 4.31 | 4.16 | 4.32 | 4.37 | 4.02 | 2.66 | 2.33 | 2.78 | 2.80 | 2.90 | 2.85 | 2.84 | 2.95 | 2.92 | 2.88 | 2.89 |
| U.comV | 4.56 | 4.39 | 4.49 | 0.00 | 1.90 | 2.03 | 2.16 | 1.84 | 1.81 | 4.51 | 4.81 | 4.08 | 4.72 | 4.60 | 4.46 | 4.35 | 4.47 | 4.36 | 4.42 | 4.41 |
| C.uniV | 4.24 | 4.25 | 4.31 | 1.90 | 0.00 | 1.14 | 1.64 | 1.18 | 0.98 | 4.26 | 4.92 | 4.44 | 4.66 | 4.63 | 4.48 | 4.40 | 4.52 | 4.52 | 4.45 | 4.45 |
| T.genIDV<br>5269V | 4.49 | 4.37 | 4.16 | 2.03 | 1.14 | 0.00 | 1.75 | 1.24 | 1.06 | 4.10 | 4.79 | 4.21 | 4.47 | 4.47 | 4.34 | 4.28 | 4.35 | 4.28 | 4.28 | 4.30 |
| M.pseV | 4.30 | 4.29 | 4.32 | 2.16 | 1.64 | 1.75 | 0.00 | 1.44 | 1.53 | 4.67 | 4.62 | 4.60 | 4.79 | 4.84 | 4.69 | 4.57 | 4.73 | 4.60 | 4.70 | 4.67 |
| A.yagV | 4.35 | 4.59 | 4.37 | 1.84 | 1.18 | 1.24 | 1.44 | 0.00 | 0.95 | 4.36 | 4.33 | 4.10 | 4.30 | 4.45 | 4.18 | 4.17 | 4.21 | 4.23 | 4.15 | 4.17 |
| L.minV | 4.21 | 4.30 | 4.02 | 1.81 | 0.98 | 1.06 | 1.53 | 0.95 | 0.00 | 4.07 | 4.51 | 4.04 | 4.30 | 4.33 | 4.11 | 4.07 | 4.12 | 4.13 | 4.01 | 4.03 |
| Foameso1 | 2.39 | 2.83 | 2.66 | 4.51 | 4.26 | 4.10 | 4.67 | 4.36 | 4.07 | 0.00 | 2.84 | 2.43 | 2.49 | 2.52 | 2.49 | 2.48 | 2.55 | 2.57 | 2.54 | 2.52 |
| Immeso1 | 2.41 | 3.28 | 2.33 | 4.81 | 4.92 | 4.79 | 4.62 | 4.33 | 4.51 | 2.84 | 0.00 | 2.99 | 2.87 | 3.00 | 2.87 | 2.89 | 2.91 | 2.85 | 2.85 | 2.86 |
| YCV | 2.23 | 3.04 | 2.78 | 4.08 | 4.44 | 4.21 | 4.60 | 4.10 | 4.04 | 2.43 | 2.99 | 0.00 | 0.95 | 0.91 | 0.93 | 0.94 | 0.97 | 0.93 | 0.93 | 0.94 |
| OFAV | 2.25 | 3.18 | 2.80 | 4.72 | 4.66 | 4.47 | 4.79 | 4.30 | 4.30 | 2.49 | 2.87 | 0.95 | 0.00 | 0.45 | 0.48 | 0.48 | 0.49 | 0.48 | 0.48 | 0.48 |
| KADV | 2.31 | 3.06 | 2.90 | 4.60 | 4.63 | 4.47 | 4.84 | 4.45 | 4.33 | 2.52 | 3.00 | 0.91 | 0.45 | 0.00 | 0.40 | 0.40 | 0.41 | 0.41 | 0.39 | 0.39 |
| DKNV | 2.28 | 2.94 | 2.85 | 4.46 | 4.48 | 4.34 | 4.69 | 4.18 | 4.11 | 2.49 | 2.87 | 0.93 | 0.48 | 0.40 | 0.00 | 0.06 | 0.13 | 0.17 | 0.11 | 0.11 |
| KSaV | 2.29 | 2.94 | 2.84 | 4.35 | 4.40 | 4.28 | 4.57 | 4.17 | 4.07 | 2.48 | 2.89 | 0.94 | 0.48 | 0.40 | 0.06 | 0.00 | 0.12 | 0.16 | 0.10 | 0.10 |
| DKV | 2.35 | 2.96 | 2.95 | 4.47 | 4.52 | 4.35 | 4.73 | 4.21 | 4.12 | 2.55 | 2.91 | 0.97 | 0.49 | 0.41 | 0.13 | 0.12 | 0.00 | 0.14 | 0.06 | 0.06 |
| CASV | 2.32 | 2.97 | 2.92 | 4.36 | 4.52 | 4.28 | 4.60 | 4.23 | 4.13 | 2.57 | 2.85 | 0.93 | 0.48 | 0.41 | 0.17 | 0.16 | 0.14 | 0.00 | 0.13 | 0.13 |
| CavV | 2.29 | 2.96 | 2.88 | 4.42 | 4.45 | 4.28 | 4.70 | 4.15 | 4.01 | 2.54 | 2.85 | 0.93 | 0.48 | 0.39 | 0.11 | 0.10 | 0.06 | 0.13 | 0.00 | 0.03 |
| NDiV | 2.28 | 2.95 | 2.89 | 4.41 | 4.45 | 4.30 | 4.67 | 4.17 | 4.03 | 2.52 | 2.86 | 0.94 | 0.48 | 0.39 | 0.11 | 0.10 | 0.06 | 0.13 | 0.03 | 0.00 |

**Extended Data Table 5.** Pairwise evolutionary distance (PED) values using RdRp alone.

|  | O.insV | X.desV | S.hybV | S.gigV | N.varV | P.opiV | U.comV | H.troV | T.genID<br>V6716V | C.uniV | T.genID<br>V5269V | M.tokV | M.pseV | A.yagV | A.stuV | E.morV | L.minV | H.mitV | Foam<br>eso1 | ACMV | BtaV | BSMV | BINV | NDiV | OFAV | KSaV | YCV |
| --- | --- | --- | --- | --- | --- | --- | --- | --- | --- | --- | --- | --- | --- | --- | --- | --- | --- | --- | --- | --- | --- | --- | --- | --- | --- | --- | --- |
| O.insV | 0.00 | 2.13 | 1.46 | 3.40 | 2.61 | 3.02 | 3.30 | 2.66 | 3.38 | 2.97 | 3.08 | 2.80 | 2.86 | 3.22 | 3.14 | 3.21 | 3.10 | 2.90 | 1.27 | 0.85 | 1.18 | 2.62 | 3.22 | 1.23 | 1.28 | 1.35 | 1.05 |
| X.desV | 2.13 | 0.00 | 2.75 | 3.28 | 2.72 | 3.05 | 3.40 | 2.96 | 2.85 | 2.90 | 2.72 | 2.88 | 2.94 | 3.38 | 2.90 | 3.16 | 2.91 | 2.94 | 2.07 | 1.95 | 2.01 | 2.94 | 3.25 | 2.21 | 2.67 | 2.41 | 2.08 |
| S.hybV | 1.46 | 2.75 | 0.00 | 3.66 | 2.70 | 3.39 | 3.59 | 3.16 | 3.22 | 3.08 | 3.13 | 2.88 | 3.02 | 3.41 | 3.19 | 2.95 | 3.14 | 2.87 | 1.55 | 1.35 | 1.71 | 2.38 | 3.31 | 1.73 | 1.78 | 1.86 | 1.55 |
| S.gigV | 3.40 | 3.28 | 3.66 | 0.00 | 3.01 | 2.97 | 2.75 | 3.27 | 3.29 | 3.21 | 3.03 | 3.25 | 3.32 | 3.03 | 3.03 | 3.20 | 2.84 | 3.27 | 3.31 | 3.01 | 3.31 | 2.42 | 2.71 | 3.35 | 2.92 | 3.30 | 3.19 |
| N.varV | 2.61 | 2.72 | 2.70 | 3.01 | 0.00 | 1.09 | 1.29 | 0.85 | 1.17 | 1.00 | 0.94 | 1.83 | 1.42 | 1.17 | 0.91 | 1.06 | 0.99 | 1.10 | 2.33 | 2.45 | 2.38 | 2.58 | 2.70 | 2.42 | 2.72 | 2.60 | 2.56 |
| P.opiV | 3.02 | 3.05 | 3.39 | 2.97 | 1.09 | 0.00 | 1.30 | 0.92 | 0.85 | 0.61 | 0.56 | 2.02 | 1.15 | 0.71 | 0.48 | 0.60 | 0.49 | 0.91 | 2.94 | 2.92 | 2.81 | 2.80 | 3.08 | 2.91 | 3.40 | 3.41 | 2.96 |
| U.comV | 3.30 | 3.40 | 3.59 | 2.75 | 1.29 | 1.30 | 0.00 | 1.39 | 1.51 | 1.35 | 1.38 | 1.94 | 1.70 | 1.27 | 1.40 | 1.27 | 1.27 | 1.57 | 3.34 | 3.10 | 2.97 | 2.68 | 2.99 | 2.96 | 3.33 | 3.28 | 2.87 |
| H.troV | 2.66 | 2.96 | 3.16 | 3.27 | 0.85 | 0.92 | 1.39 | 0.00 | 1.19 | 0.90 | 0.84 | 1.89 | 1.38 | 1.04 | 0.87 | 0.94 | 0.91 | 1.12 | 2.69 | 2.84 | 2.63 | 2.77 | 2.79 | 2.55 | 2.74 | 2.89 | 2.82 |
| T.genID<br>V6716V | 3.38 | 2.85 | 3.22 | 3.29 | 1.17 | 0.85 | 1.51 | 1.19 | 0.00 | 0.85 | 0.67 | 2.41 | 1.38 | 0.95 | 0.67 | 0.82 | 0.78 | 1.22 | 3.15 | 3.15 | 2.84 | 3.00 | 3.23 | 2.95 | 3.53 | 3.66 | 2.92 |
| C.uniV | 2.97 | 2.90 | 3.08 | 3.21 | 1.00 | 0.61 | 1.35 | 0.90 | 0.85 | 0.00 | 0.57 | 2.11 | 1.13 | 0.77 | 0.55 | 0.21 | 0.48 | 0.80 | 2.97 | 2.85 | 2.78 | 2.80 | 3.00 | 2.79 | 3.11 | 3.15 | 2.82 |
| T.genID<br>V5269V | 3.08 | 2.72 | 3.13 | 3.03 | 0.94 | 0.56 | 1.38 | 0.84 | 0.67 | 0.57 | 0.00 | 1.95 | 1.18 | 0.71 | 0.28 | 0.58 | 0.49 | 0.86 | 2.93 | 2.69 | 2.62 | 3.00 | 3.40 | 2.66 | 3.13 | 3.04 | 2.66 |
| M.tokV | 2.80 | 2.88 | 2.88 | 3.25 | 1.83 | 2.02 | 1.94 | 1.89 | 2.41 | 2.11 | 1.95 | 0.00 | 2.50 | 2.13 | 1.99 | 2.13 | 2.03 | 2.09 | 2.62 | 2.73 | 2.31 | 2.42 | 2.94 | 2.38 | 2.68 | 2.61 | 2.48 |
| M.pseV | 2.86 | 2.94 | 3.02 | 3.32 | 1.42 | 1.15 | 1.70 | 1.38 | 1.38 | 1.13 | 1.18 | 2.50 | 0.00 | 1.08 | 1.12 | 0.99 | 1.01 | 1.19 | 3.02 | 2.75 | 3.01 | 2.97 | 2.80 | 3.06 | 3.11 | 3.15 | 2.96 |
| A.yagV | 3.22 | 3.38 | 3.41 | 3.03 | 1.17 | 0.71 | 1.27 | 1.04 | 0.95 | 0.77 | 0.71 | 2.13 | 1.08 | 0.00 | 0.69 | 0.66 | 0.62 | 1.08 | 3.43 | 2.85 | 2.88 | 2.85 | 3.05 | 2.97 | 3.19 | 3.29 | 2.85 |
| A.stuV | 3.14 | 2.90 | 3.19 | 3.03 | 0.91 | 0.48 | 1.40 | 0.87 | 0.67 | 0.55 | 0.28 | 1.99 | 1.12 | 0.69 | 0.00 | 0.53 | 0.45 | 0.84 | 2.80 | 2.77 | 2.70 | 2.80 | 3.08 | 2.67 | 3.04 | 3.13 | 2.70 |
| E.morV | 3.21 | 3.16 | 2.95 | 3.20 | 1.06 | 0.60 | 1.27 | 0.94 | 0.82 | 0.21 | 0.58 | 2.13 | 0.99 | 0.66 | 0.53 | 0.00 | 0.50 | 0.77 | 3.12 | 3.09 | 3.04 | 2.51 | 2.81 | 3.02 | 3.08 | 3.04 | 2.92 |
| L.minV | 3.10 | 2.91 | 3.14 | 2.84 | 0.99 | 0.49 | 1.27 | 0.91 | 0.78 | 0.48 | 0.49 | 2.03 | 1.01 | 0.62 | 0.45 | 0.50 | 0.00 | 0.77 | 3.05 | 2.76 | 2.73 | 2.96 | 3.15 | 2.79 | 3.16 | 3.24 | 3.00 |
| H.mitV | 2.90 | 2.94 | 2.87 | 3.27 | 1.10 | 0.91 | 1.57 | 1.12 | 1.22 | 0.80 | 0.86 | 2.09 | 1.19 | 1.08 | 0.84 | 0.77 | 0.77 | 0.00 | 2.94 | 3.01 | 2.65 | 2.81 | 2.84 | 2.69 | 2.87 | 3.05 | 2.54 |
| Foameso1 | 1.27 | 2.07 | 1.55 | 3.31 | 2.33 | 2.94 | 3.34 | 2.69 | 3.15 | 2.97 | 2.93 | 2.62 | 3.02 | 3.43 | 2.80 | 3.12 | 3.05 | 2.94 | 0.00 | 1.27 | 1.34 | 2.52 | 2.87 | 1.42 | 1.52 | 1.49 | 1.27 |
| ACMV | 0.85 | 1.95 | 1.35 | 3.01 | 2.45 | 2.92 | 3.10 | 2.84 | 3.15 | 2.85 | 2.69 | 2.73 | 2.75 | 2.85 | 2.77 | 3.09 | 2.76 | 3.01 | 1.27 | 0.00 | 1.17 | 2.38 | 3.00 | 1.20 | 1.31 | 1.32 | 1.16 |
| BtaV | 1.18 | 2.01 | 1.71 | 3.31 | 2.38 | 2.81 | 2.97 | 2.63 | 2.84 | 2.78 | 2.62 | 2.31 | 3.01 | 2.88 | 2.70 | 3.04 | 2.73 | 2.65 | 1.34 | 1.17 | 0.00 | 2.69 | 3.45 | 0.11 | 0.49 | 0.07 | 0.53 |
| BSMV | 2.62 | 2.94 | 2.38 | 2.42 | 2.58 | 2.80 | 2.68 | 2.77 | 3.00 | 2.80 | 3.00 | 2.42 | 2.97 | 2.85 | 2.80 | 2.51 | 2.96 | 2.81 | 2.52 | 2.38 | 2.69 | 0.00 | 1.73 | 2.72 | 3.07 | 2.76 | 2.35 |
| BINV | 3.22 | 3.25 | 3.31 | 2.71 | 2.70 | 3.08 | 2.99 | 2.79 | 3.23 | 3.00 | 3.40 | 2.94 | 2.80 | 3.05 | 3.08 | 2.81 | 3.15 | 2.84 | 2.87 | 3.00 | 3.45 | 1.73 | 0.00 | 3.43 | 4.19 | 3.97 | 3.26 |
| NDiV | 1.23 | 2.21 | 1.73 | 3.35 | 2.42 | 2.91 | 2.96 | 2.55 | 2.95 | 2.79 | 2.66 | 2.38 | 3.06 | 2.97 | 2.67 | 3.02 | 2.79 | 2.69 | 1.42 | 1.20 | 0.11 | 2.72 | 3.43 | 0.00 | 0.49 | 0.15 | 0.56 |
| OFAV | 1.28 | 2.67 | 1.78 | 2.92 | 2.72 | 3.40 | 3.33 | 2.74 | 3.53 | 3.11 | 3.13 | 2.68 | 3.11 | 3.19 | 3.04 | 3.08 | 3.16 | 2.87 | 1.52 | 1.31 | 0.49 | 3.07 | 4.19 | 0.49 | 0.00 | 0.51 | 0.63 |
| KSaV | 1.35 | 2.41 | 1.86 | 3.30 | 2.60 | 3.41 | 3.28 | 2.89 | 3.66 | 3.15 | 3.04 | 2.61 | 3.15 | 3.29 | 3.13 | 3.04 | 3.24 | 3.05 | 1.49 | 1.32 | 0.07 | 2.76 | 3.97 | 0.15 | 0.51 | 0.00 | 0.62 |
| YCV | 1.05 | 2.08 | 1.55 | 3.19 | 2.56 | 2.96 | 2.87 | 2.82 | 2.92 | 2.82 | 2.66 | 2.48 | 2.96 | 2.85 | 2.70 | 2.92 | 3.00 | 2.54 | 1.27 | 1.16 | 0.53 | 2.35 | 3.26 | 0.56 | 0.63 | 0.62 | 0.00 |

**Extended Data Table 6.** This table lists the abbreviations for the virus names used in this article.

| **Virus name** | **Abbreviated** |
| --- | --- |
| *Yellow_head_virus* | YHV |
| *Macrobrachium rosenbergii Golda virus* | MrGV |
| *Gill-associated virus* | GAV |
| *Insect mesonivirus 1* | Immeso1 |
| *Frankliniella occidentalis associated mesonivirus 1* | Foameso1 |
| *Chinook salmon bafinivirus* | CSBV |
| *Karang Sari virus* | KSaV |
| *Dak Nong virus* | DKNV |
| *Botrylloides leachii nidovirus* | BINV |
| *Nam Dinh virus* | NDiV |
| *Cavally virus* | CavV |
| *Casuarina virus* | CASV |
| *Dianke virus* | DKV |
| *Kadiweu virus* | KADV |
| *Ofaie virus* | OFAV |
| *Yichang virus* | YCV |
| *Bontang virus* | BtaV |
| *Bradybaena similaris medionivirus* | BSMV |
| *Aphis citricidus meson-like virus* | ACMV |

## Notes

### Competing Interest Statement

The authors have declared no competing interest.

